# PBP4 is required for serum-induced cell wall thickening and antibiotic tolerance in *Staphylococcus aureus*

**DOI:** 10.1101/2024.06.19.599726

**Authors:** Elizabeth V. K. Ledger, Ruth C. Massey

## Abstract

The bacterial pathogen *Staphylococcus aureus* responds to the host environment by synthesising a thick peptidoglycan cell wall which protects the bacterium from membrane-targeting antimicrobials and the immune response. However, the proteins required for this response were previously unknown. Here, we demonstrate by three independent approaches that the penicillin binding protein PBP4 is crucial for serum-induced cell wall thickening. Firstly, mutants lacking various non-essential cell wall synthesis enzymes were tested, revealing that a mutant lacking *pbp4* was unable to generate a thick cell wall in serum. This resulted in reduced serum-induced tolerance of the *pbp4* mutant towards the last resort antibiotic daptomycin relative to wildtype cells. Secondly, we found that serum-induced cell wall thickening occurred in each of a panel of 134 clinical bacteraemia isolates, except for one strain with a naturally-occurring mutation that confers a S140R substitution in the active site of PBP4. Finally, inhibition of PBP4 with cefoxitin prevented serum-induced cell wall thickening and the resulting antibiotic tolerance in the USA300 strain and in clinical MRSA isolates. Together, this provides a rationale for combining daptomycin with cefoxitin, a PBP4 inhibitor, to potentially improve treatment outcomes for patients with invasive MRSA infections.

## Introduction

*Staphylococcus aureus* is a leading cause of invasive infections, resulting in over 12,000 cases of bacteraemia each year in the UK^1^. Methicillin susceptible *S. aureus* (MSSA) infections are often treated with front-line β-lactams such as oxacillin, however, treatment of invasive infections caused by methicillin resistant *S. aureus* (MRSA) is challenging, with patients suffering from high rates of relapse and chronic infections^2,3^. Treatment options include vancomycin and daptomycin, a cyclic lipopeptide antibiotic first approved to treat MRSA bacteraemia in 2003^4–6^. In its calcium-bound, active form it binds to the lipid phosphatidylglycerol, permeabilising the bacterial membrane^4,7,8^. This leads to a loss of intracellular ions and ATP, as well as disruption to the cell wall biosynthetic and division machinery^4,9–11^. *In vitro*, daptomycin is rapidly bactericidal but this is not the case in patients, where despite giving high doses of daptomycin intravenously, it can take several days to sterilise the bloodstream^12,13^. This slow rate of clearance allows *S. aureus* to disseminate around the body and leads to the development of secondary infections such as infective endocarditis, osteomyelitis and deep tissue abscesses^14^. As a result, patients require long hospital stays and suffer from high mortality rates^15,16^. Specifically, daptomycin fails to cure 20 – 30 % cases of MRSA bacteraemia and is associated with a 10 – 20 % mortality rate^12,13,17,18^. It is therefore important to understand why daptomycin treatment failure occurs so that new approaches can be developed to improve patient outcomes.

One explanation for daptomycin treatment failure is that the host environment induces daptomycin tolerance^19,20^. When *S. aureus* is incubated in human serum, the antimicrobial peptide LL-37 activates the GraRS two component system^19^. This leads to a reduction in the level of peptidoglycan-hydrolysing autolysins present in the cell, enabling the cell wall to thicken^19^. Since daptomycin must pass through the cell wall to reach its site of action, this thickened cell wall leads to a high level of daptomycin tolerance^19^. However, while the reduction in autolysins enables the cell wall to remain thick, it is currently unknown what proteins are required to synthesise the cell wall in serum. Understanding this is crucial so that combination therapeutic approaches can be developed to prevent daptomycin tolerance and improve patient outcomes.

## Results

### Non-essential cell wall biosynthesis genes are required for serum-induced cell wall thickening and daptomycin tolerance

The first aim of this study was to establish which *S. aureus* genes were required for cell wall thickening in human serum. As many proteins responsible for cell wall synthesis are essential and cannot be deleted, we used two mutant strains, each deficient in several non-essential enzymes involved in cell wall synthesis^21^. These strains are known as the MIN mutants as they contain the minimum peptidoglycan machinery required for peptidoglycan synthesis^21^. They have a normal growth rate and cell morphology but reduced peptidoglycan crosslinking and virulence^21^. These mutants were constructed in two genetic backgrounds, Col and Newman, and each one lacks several genes, those encoding the monofunctional transglycosylases, SgtA and SgtB, the non-essential penicillin binding proteins (PBPs), PBP3 and PBP4 and the methicillin resistance factors, FmtA and FmtB^21^. In addition, *mecA* was deleted to construct Col MIN^21^.

Firstly, we determined whether serum induced increased peptidoglycan accumulation in these strains using the D-alanine analogue HADA. This analogue is incorporated specifically into the stem peptides of the peptidoglycan and so provides a direct measurement of the amount of *de novo* peptidoglycan synthesis which occurs during the incubation period^19,22,23^. As expected from previous work^19^, wildtype (WT) bacteria grown to mid-exponential phase in tryptic soy broth (TSB-grown) had a relatively thin wall, whilst bacteria which had been grown to exponential phase and then incubated for 16 h in human serum (serum-incubated) had a significantly thicker cell wall, as shown by accumulation of HADA (Fig. 1A – B). By contrast, the MIN strains demonstrated a significantly lower level of serum-induced peptidoglycan accumulation, indicating that non-essential cell wall synthetic machinery is required for this phenotype (Fig. 1A – B).

**Figure 1.**
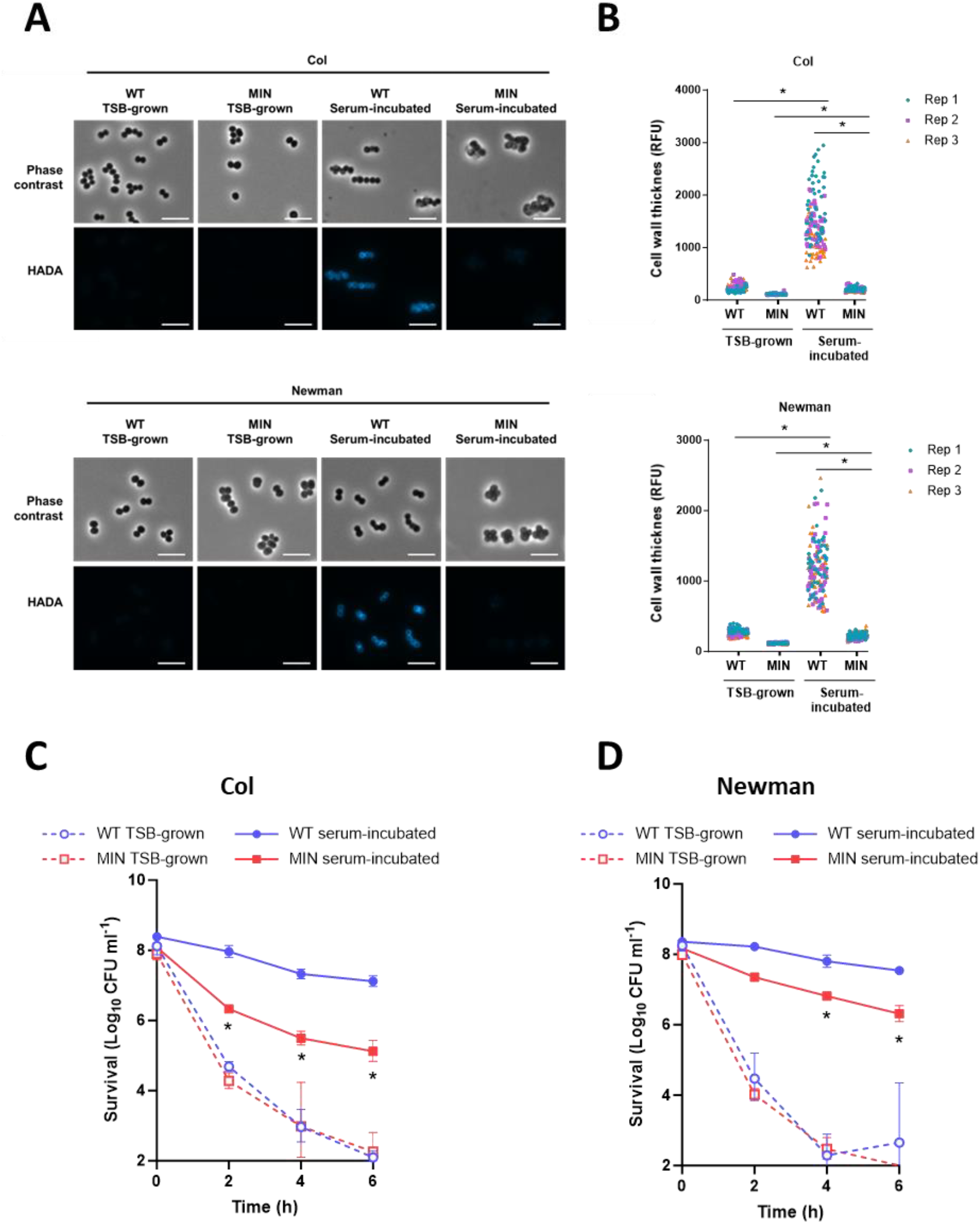
Non-essential cell wall biosynthetic proteins are required for serum-induced cell wall thickening and daptomycin tolerance. (A) TSB-grown and serum-incubated cultures of Col WT and MIN and Newman WT and MIN were generated in the presence of HADA and *de novo* cell wall synthesis determined by fluorescence microscopy. Scale bars, 5 µm. (B) Fluorescence at the peripheral cell wall of 50 cells per condition per replicate of panel A was determined. TSB-grown and serum-incubated cultures of (C) Col WT and MIN and (D) Newman WT and MIN were exposed to 80 µg ml^-1^ daptomycin for 6 h and survival determined by CFU counts. Data in B represent the fluorescence of 50 cells per replicate (150 cells total per condition). *, P ≤ 0.05 as determined by two-way ANOVA with Tukey’s *post-hoc* test. Data in C and D represent the geometric mean ± geometric standard deviation of three independent replicates. *, P ≤ 0.05 as determined by two-way ANOVA with Tukey’s *post-hoc* test; Serum-incubated WT vs MIN at indicated time-points.

To understand if the reduction in serum-induced peptidoglycan accumulation in the MIN strains affected antibiotic tolerance, we compared bacterial survival during daptomycin exposure for TSB-grown and serum-incubated cells. There were no differences in the killing kinetics of the WT and mutant strains when TSB-grown, demonstrating that the loss of non-essential cell wall synthetic enzymes did not affect daptomycin susceptibility (Fig. 1C – D). By contrast, both MIN strains were killed significantly more by daptomycin than the WT strains after serum-incubation (Fig. 1C – D). Therefore, at least one of the enzymes lacked by the MIN strains was required for serum-induced peptidoglycan accumulation and daptomycin tolerance in serum.

### PBP4 is required for serum-induced cell wall thickening and tolerance towards daptomycin

As Col MIN lacked seven enzymes relative to wild type cells and Newman MIN lacked six^21^, the next objective was to determine which of these enzymes was required for tolerance. To do this, mutants lacking each of the individual enzymes that had been deleted in the MIN strains were obtained from the Nebraska transposon mutant library (NTML)^24^, except *fmtB*, for which a mutant could not be identified, and *mecA* as this was only present in Col WT and not in Newman WT. Each of these mutants was screened for its ability to accumulate peptidoglycan during incubation in serum. Single mutants in *sgtA, sgtB, pbp3* and *fmtA* each incorporated the same amount of HADA as the WT strain during a 16 h incubation in human serum (Fig. 2A). By contrast, the mutant defective for *pbp4* incorporated significantly less HADA than WT during serum incubation (Fig. 2A). Concordantly, the *pbp4*::Tn mutant showed a 20-fold reduction in serum-induced tolerance to daptomycin compared to the WT after 6 h exposure to the antibiotic (Fig. 2B). Complementation of the *pbp4*::Tn mutant with *pbp4* expressed from its native promoter on a multi-copy plasmid restored the ability of the strain to accumulate peptidoglycan during the incubation in serum, whereas complementation with an empty plasmid did not (Fig. 2C). Complementation of the *pbp4*::Tn mutant with a functional copy of the gene also restored the serum-induced daptomycin tolerance phenotype, with the complemented strain surviving daptomycin exposure to a similar degree to the WT strain and 2-logs higher than the empty vector control (Fig. 2D). Therefore, PBP4 is required for peptidoglycan accumulation in serum and the resulting daptomycin tolerance.

**Figure 2.**
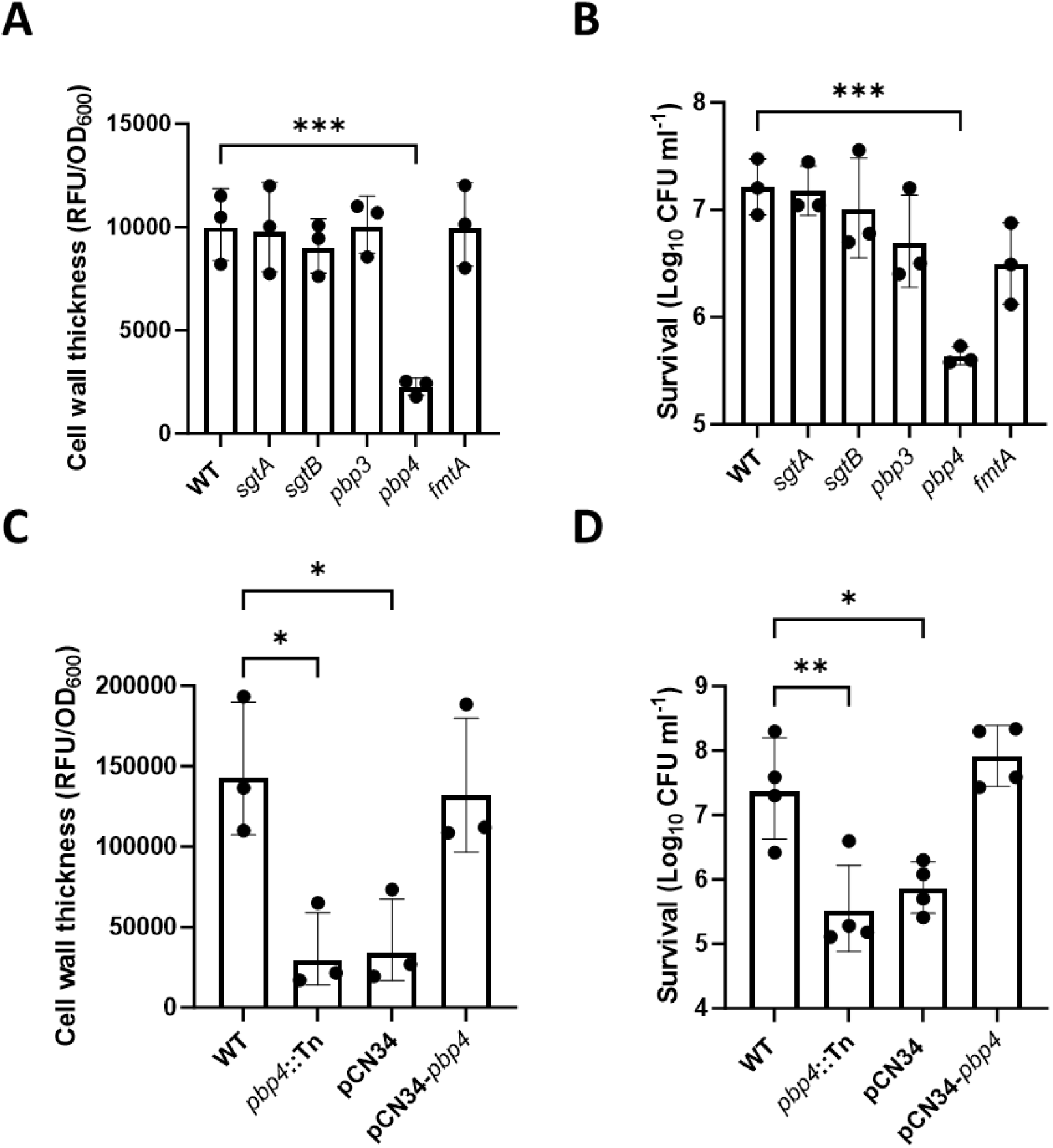
PBP4 is required for serum-induced cell wall thickening and daptomycin tolerance. (A) Cell wall thickening measured by HADA fluorescence (RFU) divided by OD_600_ of serum-incubated cultures of JE2 WT and indicated NARSA transposon mutants. (B) Survival presented as log_10_ CFU ml^-1^ of serum-incubated cultures of JE2 WT and indicated NARSA transposon mutants after a 6 h exposure to 80 µg ml^-1^ daptomycin. (C) Cell wall thickening measured by HADA fluorescence (RFU) divided by OD_600_ of serum-incubated cultures of JE2 WT, *pbp4*::Tn mutant and *pbp4*::Tn mutant complemented with empty pCN34 or pCN34 containing a WT copy of *pbp4*. (D) Survival presented as log_10_ CFU ml^-1^ of serum-incubated cultures of JE2 WT, *pbp4*::Tn mutant and *pbp4*::Tn mutant complemented with empty pCN34 or pCN34 containing a WT copy of *pbp4* after exposure to 80 µg ml^-1^ daptomycin for 6 h. Data in A and C represent the mean ± standard deviation of three independent repeats. Data in B and D represent the geometric mean ± geometric standard deviation of at least three independent repeats. Data were analysed by one-way ANOVA with Dunnett’s *post-hoc* test. *, P ≤ 0.05, **, P ≤ 0.01, ***, P ≤ 0.0001, WT vs mutant or complemented mutant.

### Serum-induced cell wall thickening and daptomycin tolerance occur in clinical isolates and are PBP4-dependent

The next objectives were to determine whether serum-induced cell wall thickening was a conserved phenotype in clinical isolates, and if so, to elucidate the mechanism and impact on daptomycin tolerance. To do this, we measured the amount of peptidoglycan synthesised by each of a panel of 134 clinical bacteraemia isolates from clonal complex 22 (CC22) during a 16 h incubation in human serum (Table S1). CC22 is highly prevalent in the UK, causing over 50 % of cases of MRSA bacteraemia^25^. Although the degree of HADA fluorescence of the strains varied after incubation in serum, it was generally high, with the exception of one isolate (ASARM124; indicated with an arrow), which showed significantly reduced fluorescence compared to the others (Fig. 3A).

**Figure 3.**
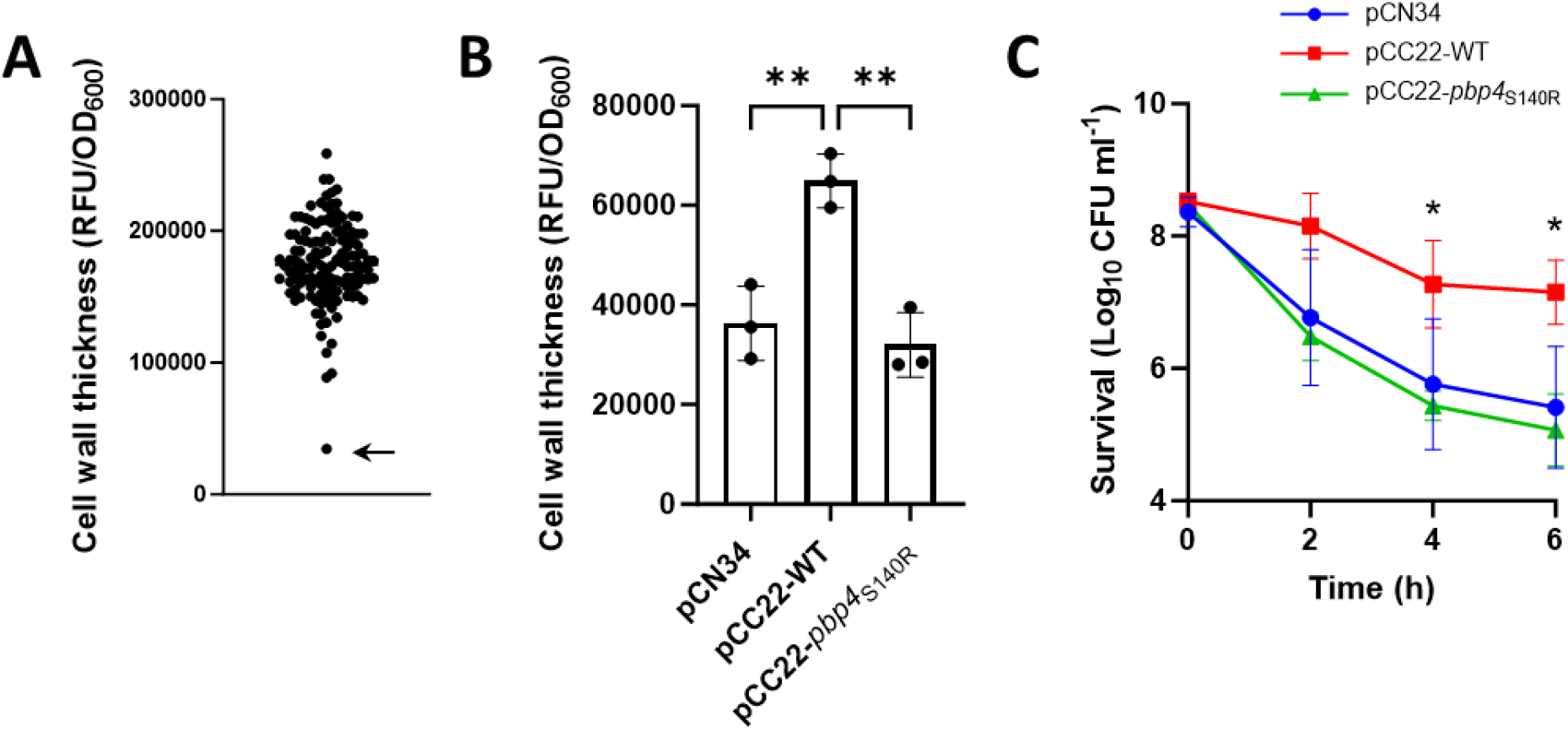
Serum-induced cell wall thickening and daptomycin tolerance occur in clinical isolates and are PBP4-dependent. (A) Cell wall thickening measured by HADA fluorescence (RFU) divided by OD_600_ of serum-incubated cultures of a panel of 134 clinical bacteraemia isolates. Each data point represents the mean of three independent repeats of one clinical isolate. The arrow indicates the strain containing the nonsynonymous mutation in *pbp4*. (B) Cell wall thickening measured by HADA fluorescence (RFU) divided by OD_600_ of serum-incubated cultures of the JE2 *pbp4*::Tn mutant complemented with empty pCN34, pCN34 containing a WT copy of *pbp4* (pCC22-WT) or pCN34 containing a copy of *pbp4* containing the SNP (pCC22-*pbp4*_S140R_). (C) Survival presented as log_10_ CFU ml^-1^ of serum-incubated cultures of the JE2 *pbp4*::Tn mutant complemented with empty pCN34, pCN34 containing a WT copy of *pbp4* (pCC22-WT) or pCN34 containing a copy of *pbp4* containing the point mutation (pCC22-*pbp4* _S140R_) after a 6 h exposure to 80 µg ml^-1^ daptomycin. Data in B represent the mean ± standard deviation of three independent repeats and data in C represent the geometric mean ± geometric standard deviation of three independent repeats. Data in B were analysed by one-way ANOVA with Dunnett’s *post-hoc* test (**, P ≤ 0.01) and data in C were analysed by two-way ANOVA with Dunnett’s *post-hoc* test (*, P ≤ 0.05; pCN34 vs pCC22-WT at indicated time-points).

The next objective was to understand why this strain did not respond to serum by thickening its cell wall. To do this, we compared the sequences of the *pbp4* genes encoded by each of the clinical isolates. This revealed that the amino acid sequences of PBP4 in each of these strains were identical, with the exception of ASARM124, the strain with a very low level of HADA incorporation. This isolate contained a nonsynonymous mutation at position 420 of *pbp4* (A>C), resulting in a S140R substitution. Serine 140 is in the active site of the enzyme and so it is likely that this disrupts its catalytic activity^26^. To confirm that this variant of PBP4 was less active, the *pbp4*::Tn mutant from the NTML was complemented with either the WT sequence of *pbp4* present in the other isolates (pCC22-WT) or with the variant (pCC22-*pbp4*_S140R_). The ability of these strains to incorporate HADA during incubation in serum and become tolerant towards daptomycin was then measured. As expected, complementation with the WT copy of *pbp4* led to significantly higher levels of HADA incorporation than the empty vector control (Fig. 3B). When the *pbp4*::Tn mutant was complemented with the *pbp4* variant, no increase in HADA incorporation was seen compared to the empty vector control, demonstrating that the protein variant encoded is inactive (Fig. 3B). In line with this, complementation with the WT *pbp4* sequence led to significantly higher levels of daptomycin tolerance than the empty plasmid whereas complementation with the variant *pbp4* gene did not restore tolerance (Fig. 3C). Taken together, these experiments showed that peptidoglycan accumulation occurs when clinical isolates are incubated in serum and that this is dependent on PBP4.

### Inhibition of PBP4 with cefoxitin reduces serum-induced cell wall thickening and daptomycin tolerance in USA300 JE2 and clinical MRSA strains

As a final examination of the role of PBP4 in serum-induced tolerance, we used cefoxitin, a second-generation cephalosporin antibiotic which inhibits all PBPs at growth-inhibitory concentrations, but due to its high affinity for PBP4, selectively inhibits this PBP at lower concentrations^27,28^. The minimum inhibitory concentration (MIC) of cefoxitin in both JE2 WT and the *pbp4*::Tn mutant was 8 µg ml^-1^. Firstly, we tested a range of concentrations of cefoxitin to determine the optimum concentration to use. JE2 WT was incubated in serum for 16 h with cefoxitin and HADA to determine a concentration which prevented cell wall thickening from occurring. Cefoxitin led to dose dependent inhibition of HADA incorporation, with a concentration of 2 µg ml^-1^ resulting in the same level of HADA fluorescence as was observed for the *pbp4*::Tn mutant (Fig. 4A). Next, we investigated whether inhibition of PBP4 with cefoxitin reduced daptomycin tolerance. JE2 WT and the *pbp4*::Tn mutant were incubated in serum for 16 h supplemented or not with 2 µg ml^-1^ cefoxitin before 80 µg ml^-1^ daptomycin was added and survival measured. At the sub-MIC concentration of cefoxitin used, there was no loss of viability of *S. aureus* in serum, with CFU counts remaining at the starting inoculum (2 × 10^8^ CFU ml^-1^) throughout the assay, as occurs in the absence of cefoxitin (Fig. 4B). Subsequent daptomycin challenge revealed that, as expected, the WT strain incubated in serum without cefoxitin had high levels of tolerance to the lipopeptide antibiotic, and this was significantly reduced in the *pbp4*::Tn mutant similarly incubated (Fig. 4C). However, WT cells incubated in serum containing cefoxitin had significantly reduced daptomycin tolerance, with bacterial killing occurring at levels similar to the *pbp4*::Tn mutant (Fig. 4C). The lack of effect of cefoxitin on the daptomycin tolerance of the *pbp4*::Tn mutant confirmed that the antibiotic acted via PBP4 and not via off-target effects.

**Figure 4.**
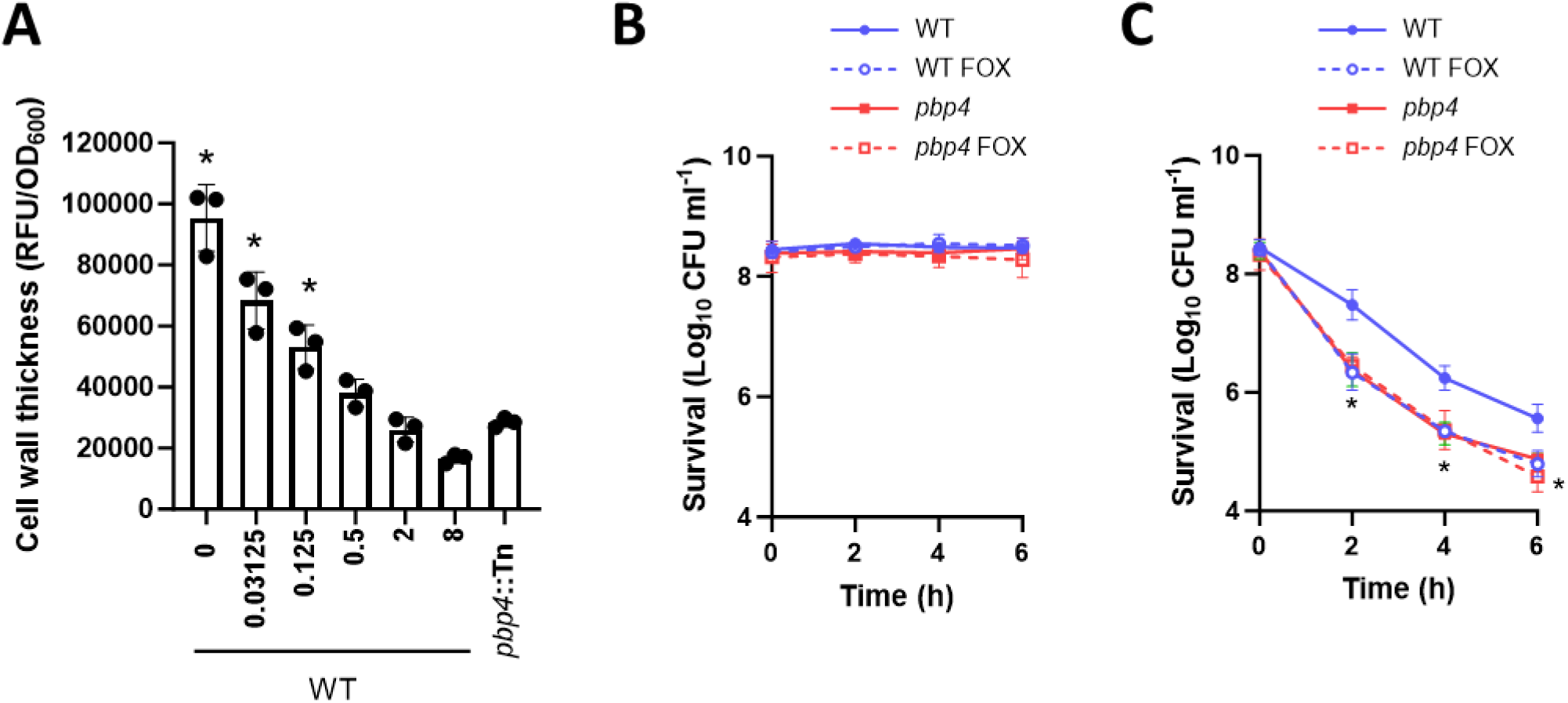
Inhibition of PBP4 with cefoxitin reduces cell wall thickening and daptomycin tolerance in JE2. (A) Cell wall thickening measured by HADA fluorescence (RFU/OD_600_) of JE2 WT after a 16 h incubation in serum supplemented with the indicated concentration of cefoxitin or the *pbp4*::Tn mutant incubated in serum alone. JE2 WT or the *pbp4*::Tn mutant were incubated in serum supplemented, or not, with 2 µg ml^-1^ cefoxitin (FOX) for 16 h before (B) log_10_ CFU ml^-1^ were determined over 6 h and (C) 80 µg ml^-1^ daptomycin was added and log_10_ CFU ml^-1^ were determined over 6 h. Data in A represent the mean ± standard deviation of three independent replicates. Data in B and C represent the geometric mean ± geometric standard deviation of three independent replicates. Data in A were analysed by one-way ANOVA with Dunnett’s *post-hoc* test (*, P ≤ 0.05; *pbp4*::Tn vs other conditions). Data in B and C were analysed by two-way ANOVA with Dunnett’s *post-hoc* test (*, P ≤ 0.05; WT vs other conditions at indicated time-points).

Next, we determined whether cefoxitin was also effective at reducing tolerance in the clinical MRSA strains. To do this, four clinical isolates were chosen which incorporated HADA into their cell walls during incubation in serum (ASARM141, ASARM179, ASARM184 and ASARM208), along with the strain containing the inactivating point mutation in *pbp4* (ASARM124). Like JE2, these strains were all resistant to cefoxitin (MICs 8 – 16 µg ml^-1^). Supplementation of the serum with cefoxitin led to dose dependent inhibition of HADA incorporation in each of the strains encoding WT PBP4 (Fig. 5A – D), whereas no significant reduction in fluorescence was observed in ASARM124 (Fig. 5E). For each of the strains with WT PBP4, 2 µg ml^-1^ cefoxitin was sufficient to reduce the fluorescence to the level of the strain lacking PBP4 (Fig. 5A – E) and so this concentration was chosen to determine whether cefoxitin was able to reduce daptomycin tolerance in these clinical strains.

**Figure 5.**
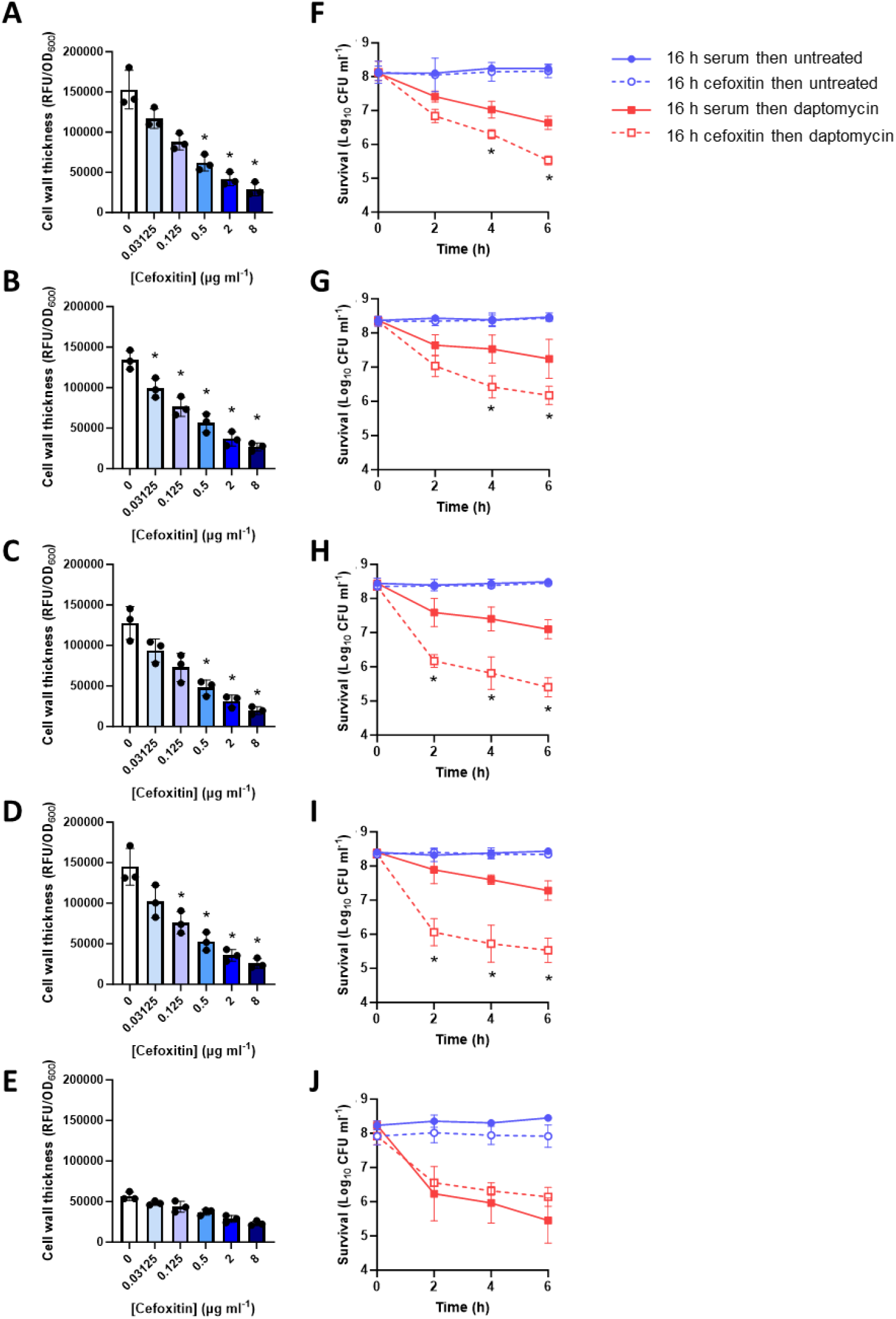
Inhibition of PBP4 with cefoxitin reduces cell wall thickening and daptomycin tolerance in clinical isolates. Cell wall thickening as measured by HADA fluorescence (RFU/OD_600_) of (A) ASARM141, (B) ASARM179, (C) ASARM184, (D) ASARM208 and (E) ASARM124 after a 16 h incubation in serum supplemented with the indicated concentration of cefoxitin. Survival as measured by log_10_ CFU ml^-1^ of (F) ASARM141, (G) ASARM179, (H) ASARM184, (I) ASARM208 and (J) ASARM124 following incubation for 16 h in serum supplemented or not with 2 µg ml^-1^ cefoxitin and the addition, or not, of 80 µg ml^-1^ daptomycin for 6 h. Data in A – E represent the mean ± standard deviation of three independent replicates. Data in F – J represent the geometric mean ± geometric standard deviation of three independent replicates. Data in A – E were analysed by one-way ANOVA with Dunnett’s *post-hoc* test (*, P ≤ 0.05; serum alone vs serum supplemented with cefoxitin). Data in F – J were analysed by two-way ANOVA with Sidak’s *post-hoc* test (*, P ≤ 0.05; daptomycin survival in serum alone vs serum supplemented with cefoxitin).

Next, each of these clinical strains was incubated for 16 h in human serum supplemented, or not, with 2 µg ml^-1^ cefoxitin before being exposed, or not, to 80 µg ml^-1^ daptomycin. As a sub-MIC concentration of cefoxitin was used, this antibiotic had no effect on bacterial survival during the 16 h incubation or in the following 6 h, with the initial inoculum of 2 × 10^8^ CFU ml^-1^ remaining at the end of the assay (Fig. 5F – J). By 6 h, addition of daptomycin resulted in approximately a 1-log reduction in CFU counts of each of the strains with WT copies of PBP4 (Fig. 5F – I). In line with the data above, the defective PBP4 in ASARM124 resulted in this strain showing reduced daptomycin tolerance compared to the other strains, with daptomycin alone causing a >2-log reduction in CFU counts (Fig. 5J). In each of the strains encoding WT PBP4, addition of cefoxitin significantly reduced daptomycin tolerance (Fig. 5F – I). However, in agreement with cefoxitin showing high selectivity for PBP4 and PBP4 being required for tolerance, addition of this antibiotic had no effect on the daptomycin tolerance of ASARM124 (Fig. 5J). Taken together, inhibition of PBP4 with cefoxitin reduces the ability of *S. aureus* to accumulate peptidoglycan in serum and reduces daptomycin tolerance.

## Discussion

*S. aureus* is a major cause of chronic and relapsing infections due to a lack of effective clearance by antibiotics and the immune system. The thickened cell wall that *S. aureus* synthesises during incubation in human serum may contribute to this, as it protects the bacterium from the last resort antibiotic daptomycin and from phagocytosis^19,29^. Inhibiting this cell wall thickening may therefore reduce antibiotic tolerance and increase clearance by immune cells. However, the proteins required for serum-induced cell wall thickening are unknown and so the aims of this work were to determine the mechanism by which the cell wall thickens and to develop a combination therapeutic approach to inhibit it. This work demonstrates that PBP4 is crucial for peptidoglycan accumulation in serum and highlights its potential as a therapeutic target the enhance the efficacy of daptomycin.

PBP4 is an uncanonical PBP, lacking the transglycosylase domain possessed by the other PBPs and having only the transpeptidase domain, with which it crosslinks the peptidoglycan units together via a pentaglycine bridge^30^. The action of PBP4 results in the *S. aureus* cell wall being highly crosslinked, with crosslinks present in 80 – 90 % of peptidoglycan units, and *pbp4* mutants having a significantly less crosslinked wall than wildtype cells^28,31^. This is thought to provide extra strength to the wall and compensate for the relatively short glycan chain length, which average 6 disaccharide units in *S. aureus* compared to up to over 500 in *Bacillus subtilis*^32,33^.

PBP4 is known to contribute to high level beta-lactam resistance in MRSA, however, it has not been linked to daptomycin susceptibility previously^34,35^. Studies investigating synergy between daptomycin and beta-lactams identified that daptomycin and cefoxitin were not synergistic and that beta-lactams need to possess PBP1-targeting activity to synergise with daptomycin^36,37^. The mechanisms behind this are unclear, however are thought to be due to beta-lactams affecting the properties of the cell envelope and/or increasing the binding of daptomycin^37–39^. However, these experiments were carried out in laboratory media and performance of antibiotics in media is not always reflective of efficacy under host conditions. Therefore, our finding that PBP4 is crucial for the development of daptomycin tolerance under host conditions raises the possibility that a PBP4 inhibitor may enhance daptomycin treatment efficacy *in vivo*.

While PBP4 is non-essential under laboratory conditions, it is important for virulence and survival inside the host, as for example a deletion mutant shows reduced virulence in a murine osteomyelitis model^40^. Clinical isolates with mutations in *pbp4* have previously been identified, however, these mutations are not inactivating mutations and are thought to enable enhanced beta-lactam resistance^41,42^. To our knowledge, this is the first clinical bacteraemia isolate identified with a deleterious mutation in *pbp4*. The importance of PBP4 for survival in the host and for beta-lactam resistance is likely to provide a strong selective pressure to retain PBP4 activity, meaning that targeting PBP4 to enhance daptomycin activity is likely to be effective in the vast majority of strains. Indeed, the finding here that serum induces cell wall thickening in over 99% of the clinical bacteraemia isolates tested demonstrates that this phenomenon is not only frequent, but also not confined to laboratory strains, occurring in strains which cause infections typically treated with daptomycin.

We have previously shown that inhibition of cell wall synthesis with fosfomycin reduces daptomycin tolerance, however, evidence that a daptomycin-fosfomycin combination is safe and effective is limited^19,20^. For example, one report demonstrated that while the combination therapy reduced rates of treatment failure compared to daptomycin monotherapy, there was no increase in patient survival rates and combination therapy was associated with a higher number of adverse events^43^. Therefore, while inhibition of cell wall synthesis is a promising strategy for enhancing daptomycin treatment efficacy, the choice of cell wall synthesis inhibitor needs to be carefully considered.

There has been much interest in combining daptomycin with a beta-lactam and this combination is synergistic *in vitro* and *in vivo*^39,44–47^. However, the mechanism behind the synergy is not completely understood and there is debate as to which beta-lactam is most effective. Additionally, in several trials, the daptomycin/beta-lactam combination was associated with a higher rate of adverse reactions than monotherapy^2,48^. However, to date there has not been a large-scale trial investigating whether beta-lactams improve daptomycin efficacy^49^. The SNAP trial is attempting to address this by recruiting 7000 patients with *S. aureus* bacteraemia to determine, amongst other treatment options, whether cefazolin increases daptomycin efficacy in MRSA bacteraemia cases^49^. However, to date there is very little data informing on which beta-lactam is most effective in combination with daptomycin at treating the infection while minimising adverse events, and no clinical trials combining daptomycin with cefoxitin have been performed.

Our data support that inhibition of PBP4, in this case with cefoxitin, was able to reduce daptomycin tolerance in each of the tested clinical MRSA strains. The concentration of cefoxitin used was below the EUCAST breakpoint concentration of 4 µg ml^-1^ and clinically-achievable^50^, demonstrating that this may be a viable therapeutic option to improve daptomycin treatment.

Finally, it has also recently been shown that the cell wall thickening triggered by incubation in serum also conceals opsonins bound to surface proteins and lipoteichoic acid, reducing phagocytosis by neutrophils^29^. Therefore, addition of cefoxitin into the treatment regimen may not only reduce daptomycin tolerance but also enhance killing by the immune system, leading to faster bacterial clearance and improved patient outcomes.

In summary, we have shown that cell wall thickening occurs in clinical bacteraemia isolates in response to human serum and that it is dependent on PBP4. This provides a rationale for combining daptomycin with a PBP4 inhibitor to reduce antibiotic tolerance and improve patient outcomes.

## Materials and methods

### Bacterial strains and growth conditions

The bacterial strains used in this study are shown in Table 1. The clinical isolates are shown in Table S1. *S. aureus* was grown in TSB for 16 h at 37 °C with shaking (180 rpm) to reach stationary phase. When required, TSB was supplemented with erythromycin (10 µg ml^-1^) or kanamycin (90 µg ml^-1^).

**Table 1.**
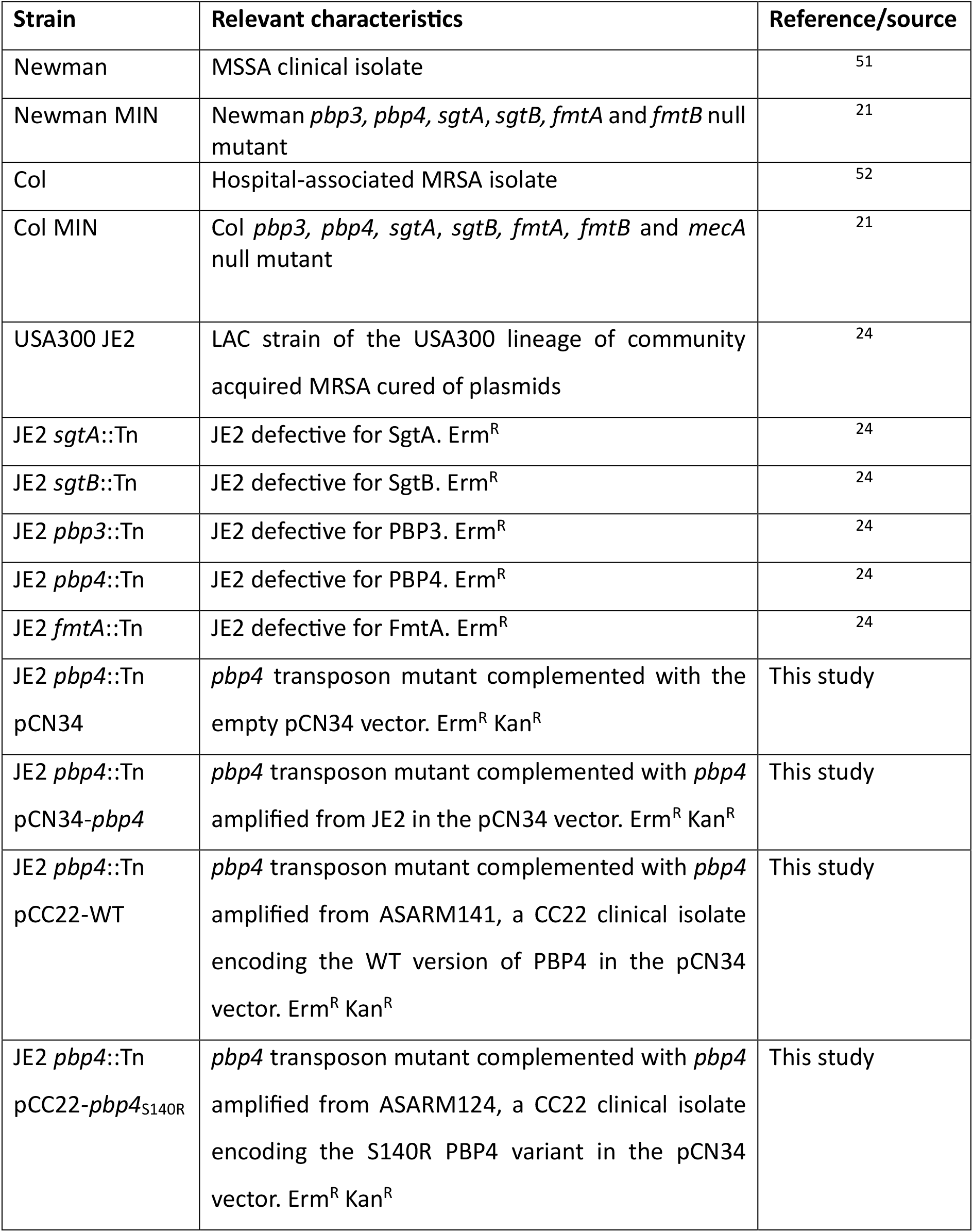
Strains used in this study.

| Strain | Relevant characteristics | Reference/source |
| --- | --- | --- |
| Newman | MSSA clinical isolate | 51 |
| Newman MIN | Newman *pbp3*, *pbp4*, *sgtA*, *sgtB*, *fmtA* and *fmtB* null mutant | 21 |
| Col | Hospital-associated MRSA isolate | 52 |
| Col MIN | Col *pbp3*, *pbp4*, *sgtA*, *sgtB*, *fmtA*, *fmtB* and *mecA* null mutant | 21 |
| USA300 JE2 | LAC strain of the USA300 lineage of community acquired MRSA cured of plasmids | 24 |
| JE2 *sgtA*::Tn | JE2 defective for SgtA. Erm<sup>R</sup> | 24 |
| JE2 *sgtB*::Tn | JE2 defective for SgtB. Erm<sup>R</sup> | 24 |
| JE2 *pbp3*::Tn | JE2 defective for PBP3. Erm<sup>R</sup> | 24 |
| JE2 *pbp4*::Tn | JE2 defective for PBP4. Erm<sup>R</sup> | 24 |
| JE2 *fmtA*::Tn | JE2 defective for FmtA. Erm<sup>R</sup> | 24 |
| JE2 *pbp4*::Tn pCN34 | *pbp4* transposon mutant complemented with the empty pCN34 vector. Erm<sup>R</sup> Kan<sup>R</sup> | This study |
| JE2 *pbp4*::Tn pCN34-*pbp4* | *pbp4* transposon mutant complemented with *pbp4* amplified from JE2 in the pCN34 vector. Erm<sup>R</sup> Kan<sup>R</sup> | This study |
| JE2 *pbp4*::Tn pCC22-WT | *pbp4* transposon mutant complemented with *pbp4* amplified from ASARM141, a CC22 clinical isolate encoding the WT version of PBP4 in the pCN34 vector. Erm<sup>R</sup> Kan<sup>R</sup> | This study |
| JE2 *pbp4*::Tn pCC22-*pbp4*<sub>S140R</sub> | *pbp4* transposon mutant complemented with *pbp4* amplified from ASARM124, a CC22 clinical isolate encoding the S140R PBP4 variant in the pCN34 vector. Erm<sup>R</sup> Kan<sup>R</sup> | This study |

To generate TSB-grown bacteria, stationary phase overnight cultures were diluted to 10^7^ CFU ml^-1^ in TSB and incubated at 37 °C with shaking (180 rpm) until 2 × 10^8^ CFU ml^-1^. To generate serum-incubated cultures, TSB-grown bacteria were centrifuged and resuspended in human serum from male AB plasma (Sigma) and incubated for 16 h at 37 °C with shaking (180 rpm). As *S. aureus* is unable to replicate in human serum, there was no change in CFU counts during this incubation in serum. Where appropriate, serum was supplemented with a sub-lethal concentration of cefoxitin (2 µg ml^-1^).

### Construction of strains

The JE2 *pbp4*::Tn mutant was complemented by cloning the *pbp4* gene along with its promoter and terminator regions into the multi-copy *E. coli/S. aureus* shuttle vector pCN34^53^. *pbp4* was amplified from JE2 using *pbp4*_Fw (5’-GATGCTTGAATCAATACGCTATGTCG -3’) and *pbp4*_Rev (5’-AGTCAGGTACCGTGTTGCCCCAAAATACCGC -3’) and from CC22 strains using *pbp4*_Fw and *pbp4*_CC22_Rev (5’-AGTCAGGTACCAGTATTGCCCCAAAATACCGTTC -3’). Fragments were then digested with EcoRI and KpnI and ligated into digested pCN34 using T4 ligase. Plasmids were transformed into *E. coli* DC10B, then electroporated into *S. aureus* RN4220 and transduced into the JE2 *pbp4*::Tn mutant using ϕ11.

### Measurements of cell wall thickening by incorporation of HADA by fluorescence microscopy

*De novo* peptidoglycan synthesis was determined by measuring incorporation of HCC-amino-D-alanine (HADA), a fluorescent D-amino acid analogue which is incorporated into the pentapeptide chain of peptidoglycan as it is synthesised^19,23^. TSB-grown cultures and serum-incubated cultures were generated as above with the addition of 25 µM HADA. Cultures were washed three times in PBS, fixed in 4 % paraformaldehyde, spotted onto agarose on microscope slides and imaged using a Zeiss Axio Imager.A1 microscope coupled to an AxioCam MRm and a 100× objective using a DAPI filter set. The fluorescence of 50 cells per replicate (150 cells total) was quantified using Zen 2012 software (blue edition).

### Measurements of cell wall thickening by incorporation of HADA by plate reader

As above, serum-incubated cultures were generated in the presence of 25 µM HADA before being washed in PBS and resuspended in PBS. Aliquots (200 µl) were transferred to black-walled 96 well plates and the fluorescence measured using a TECAN Infinite 200 PRO microplate reader (excitation 405 nm; emission 460 nm). Fluorescence values were divided by OD_600_ readings to normalise for differences in cell density between samples. For the clinical isolates, cultures were grown for 16 h in 200 µl TSB in 96 well plates, before being diluted 10-fold, incubated in serum supplemented with 25 µM HADA in 96 well plates for 16 h, washed in PBS, resuspended in PBS and fluorescence measured as above.

### Bacterial antibiotic killing assay

TSB-grown and serum-incubated cultures were exposed to 80 µg ml^-1^ daptomycin in serum for 6 h and CFU counts determined every 2 h for 6 h or at the 6 h time point only. TSB-grown cultures were centrifuged and resuspended in serum immediately before the daptomycin was added. Where appropriate, serum was supplemented with a sub-lethal concentration of cefoxitin (2 µg ml^-1^).

### Determination of antibiotic MIC

MICs were determined using the broth microdilution protocol^54^. Two-fold serial dilutions of cefoxitin were prepared in 200 µl TSB per well in a 96-well plate. Wells were inoculated at 5 × 10^5^ CFU ml^-1^ and incubated statically at 37 °C for 16 h. The MIC was the lowest concentration with no visible growth.

### Statistical analyses

CFU data were log_10_ transformed and presented as the geometric mean ± geometric standard deviation. Non-CFU data were presented as mean ± standard deviation. All experiments consisted of at least three independent biological replicates and were analysed by one-way or two-way ANOVA with appropriate *post-hoc* multiple comparisons test using GraphPad Prism (V10.0) as described in the figure legends.

## Acknowledgements

Mariana Pinho (ITQB NOVA) is thanked for providing strains. Vladimir Pelicic (Imperial College London) is thanked for providing access to the fluorescent microscope. Simon Foster (University of Sheffield) is thanked for providing HADA. All authors acknowledge the provision of strains by the Network on Antimicrobial Resistance in Staphylococcus aureus (NARSA) Program: under NIAID/ NIH Contract No. HHSN272200700055C.

