## Supplementary data for "PBP4 is required for serum-induced cell wall thickening and antibiotic tolerance in *Staphylococcus aureus*"

**Supplementary data file**

Supplementary Table 1

**Supplementary Table 1**. Details of the clinical strains used in this study.

| **Strain ID** | **Accession number** | ***pbp4* SNP** | **HADA (RFU/$OD_{600}$)** |
|---|---|---|---|
| ASARM61 | ERR084492 | No | 151274.5 |
| ASARM70 | ERR084501 | Synonymous 879G<A | 152697.6 |
| ASARM71 | ERR084502 | No | 126439.4 |
| ASARM72 | ERR084503 | No | 150334.2 |
| ASARM73 | ERR084504 | No | 111233 |
| ASARM74 | ERR084505 | No | 159019.9 |
| ASARM77 | ERR084506 | No | 142518.9 |
| ASARM76 | ERR084507 | No | 183836.5 |
| ASARM75 | ERR084508 | No | 176864.6 |
| ASARM80 | ERR084509 | No | 211497.7 |
| ASARM79 | ERR084510 | No | 164836 |
| ASARM59 | ERR084493 | No | 165364.6 |
| ASARM83 | ERR084513 | No | 147809.8 |
| ASARM84 | ERR084514 | No | 142095.5 |
| ASARM86 | ERR084516 | No | 226125 |
| ASARM87 | ERR084517 | No | 215884.5 |
| ASARM89 | ERR084519 | No | 182293.2 |
| ASARM62 | ERR084494 | No | 177857.7 |
| ASARM93 | ERR084522 | No | 172872.2 |
| ASARM95 | ERR084523 | No | 168433.9 |
| ASARM96 | ERR084524 | No | 125618.6 |
| ASARM97 | ERR084525 | No | 169614.1 |
| ASARM99 | ERR084527 | No | 165651.1 |
| ASARM100 | ERR084528 | No | 176369.9 |
| ASARM101 | ERR084529 | No | 166845 |
| ASARM102 | ERR084530 | No | 163152.8 |
| ASARM103 | ERR084531 | No | 156151.4 |
| ASARM105 | ERR084533 | No | 161566.1 |
| ASARM107 | ERR084534 | No | 175022.2 |
| ASARM109 | ERR084535 | No | 177724.2 |
| ASARM114 | ERR084540 | No | 160135.4 |
| ASARM64 | ERR109523 | No | 110420.5 |
| ASARM116 | ERR084541 | No | 164537.7 |
| ASARM117 | ERR084542 | No | 190559.9 |

| | | | |
|---|---|---|---|
| ASARM118 | ERR084543 | No | 152330.6 |
| ASARM119 | ERR084544 | No | 180603.5 |
| ASARM120 | ERR084545 | No | 160555.6 |
| ASARM121 | ERR084546 | No | 170425.6 |
| ASARM122 | ERR084547 | No | 198076.5 |
| ASARM124 | ERR084548 | Nonsynonymous 420 A<C | 31339.64 |
| ASARM125 | ERR084549 | No | 163946.5 |
| ASARM126 | ERR084550 | No | 181816.5 |
| ASARM65 | ERR084497 | No | 120655.6 |
| ASARM127 | ERR084551 | No | 133460.6 |
| ASARM128 | ERR084552 | No | 164894.7 |
| ASARM132 | ERR084554 | No | 162184.1 |
| ASARM133 | ERR084556 | No | 188048.5 |
| ASARM134 | ERR084557 | No | 157260.9 |
| ASARM135 | ERR084558 | No | 182532.5 |
| ASARM136 | ERR084559 | No | 139611.1 |
| ASARM137 | ERR084560 | No | 183746.8 |
| ASARM67 | ERR084498 | No | 186761 |
| ASARM138 | ERR084561 | No | 153196.6 |
| ASARM139 | ERR084562 | No | 178510.9 |
| ASARM140 | ERR084563 | No | 176486.3 |
| ASARM141 | ERR084564 | No | 225584.8 |
| ASARM142 | ERR084565 | No | 153257.6 |
| ASARM143 | ERR084566 | No | 211622.6 |
| ASARM144 | ERR084567 | No | 149930.1 |
| ASARM145 | ERR084568 | No | 170581.9 |
| ASARM68 | ERR084499 | No | 209713.3 |
| ASARM148 | ERR084571 | No | 188189.5 |
| ASARM154 | ERR084575 | No | 155028.3 |
| ASARM153 | ERR084576 | No | 151903.4 |
| ASARM155 | ERR084578 | No | 198692.4 |
| ASARM160 | ERR084579 | No | 183246.7 |
| ASARM69 | ERR084500 | No | 152676.9 |
| ASARM162 | ERR084581 | No | 142612.7 |
| ASARM164 | ERR084582 | No | 169513.2 |
| ASARM163 | ERR084583 | No | 156410.4 |
| ASARM166 | ERR084584 | No | 162167.1 |
| ASARM165 | ERR084585 | No | 195943 |
| ASARM167 | ERR084586 | No | 183259 |
| ASARM168 | ERR084587 | No | 161667.5 |
| ASARM169 | ERR084638 | No | 179021.9 |

| | | | |
|---|---|---|---|
| ASARM179 | ERR084648 | No | 153559.6 |
| ASARM181 | ERR084650 | No | 159738.3 |
| ASARM183 | ERR084652 | No | 227859.1 |
| ASARM184 | ERR084653 | No | 197405.9 |
| ASARM170 | ERR084639 | No | 186022.6 |
| ASARM191 | ERR084658 | No | 158991.8 |
| ASARM193 | ERR084659 | No | 211593.2 |
| ASARM199 | ERR084664 | No | 169671 |
| ASARM200 | ERR084665 | No | 160345.9 |
| ASARM201 | ERR084666 | No | 168708.7 |
| ASARM171 | ERR084640 | No | 147242.5 |
| ASARM203 | ERR084667 | No | 213237.2 |
| ASARM204 | ERR084668 | No | 164123.9 |
| ASARM205 | ERR084669 | No | 178405.3 |
| ASARM208 | ERR084671 | No | 138694.8 |
| ASARM209 | ERR084672 | No | 168964.2 |
| ASARM207 | ERR084673 | No | 143243.2 |
| ASARM211 | ERR084675 | No | 137671 |
| ASARM212 | ERR084676 | No | 131155.5 |
| ASARM172 | ERR084641 | No | 169392.5 |
| ASARMLT1 | ERR084678 | No | 166094.2 |
| ASARMLT2 | ERR084679 | No | 184064.8 |
| ASARMLT3 | ERR084680 | No | 167359 |
| ASARM195 | ERR084714 | No | 155606.3 |
| ASARM176 | ERR084645 | No | 154856.3 |
| ASARM177 | ERR084646 | No | 162984.9 |
| ASARM217 | ERR171907 | No | 155872.8 |
| ASARM220 | ERR171908 | No | 176502.9 |
| ASARM222 | ERR171910 | No | 178102.1 |
| ASARM223 | ERR171911 | No | 188774.3 |
| ASARM224 | ERR171912 | No | 133295.3 |
| ASARM110 | ERR223125 | No | 172142.3 |
| ASARM108 | ERR223118 | No | 197180.4 |
| ASASM42 | ERR109502 | No | 206715.9 |
| ASASM56 | ERR109515 | No | 148914.5 |
| ASASM12 | ERR109476 | No | 163313.1 |
| ASASM61 | ERR109520 | No | 171618.4 |
| ASASM64 | ERR084496 | No | 151796.7 |
| ASASM71 | ERR109528 | No | 183412 |
| ASASM73 | ERR109530 | No | 156821.9 |
| ASASM90 | ERR109540 | No | 164555.9 |
| ASASM96 | ERR109546 | No | 150968 |

| | | | |
|---|---|---|---|
| ASASM97 | ERR109547 | No | 197283.9 |
| ASASM120 | ERR109567 | No | 83346.79 |
| ASASM132 | ERR109578 | No | 205175.5 |
| ASASM138 | ERR109584 | No | 139711.7 |
| ASASM140 | ERR109586 | No | 152372.3 |
| ASASM125 | ERR109572 | No | 131370.1 |
| ASASM127 | ERR109574 | No | 142147.3 |
| ASASM181 | ERR109623 | No | 159693 |
| ASASM190 | ERR109630 | No | 147543 |
| ASASM246 | ERR114858 | No | 186166.2 |
| ASASM262 | ERR114873 | No | 111483.3 |
| ASASM390 | ERR172029 | No | 107684.2 |
| ASASM392 | ERR172031 | No | 173241.6 |
| ASASM430 | ERR172068 | No | 154671.3 |
| ASASM168 | ERR223120 | No | 162396.6 |
